# Understanding epigenetic regulation in non-model plants: Transcriptomic responses to seed demethylation in leaves and roots of the annual herb *Erodium cicutarium* (Geraniaceae)

**DOI:** 10.1101/2025.11.17.688907

**Authors:** Rubén Martín-Blázquez, Mónica Medrano, Conchita Alonso

**Author notes:** **Corresponding autor:** Conchita Alonso.

## Abstract

Epigenetic regulation has emerged as a significant element in adaptation to heterogeneous and stressful environments, with modifications in DNA methylation being particularly relevant in plants. DNA methylation inhibitors have been used to investigate the relationship between DNA methylation and plastic plant phenotypes. However, their effect in gene expression regulation along lifetime remains understudied in non-model plants. Here, we analyze the effects of seed exposure to 5-azacytidine (5-azaC) in plant gene regulation. Scarified seeds from a single inbred line of *Erodium cicutarium* were soaked for 48 h in either water or a low concentration solution of 5-azaC before sowing. Subsequently, RNA was extracted from juvenile roots, juvenile leaves and adult leaves, and their transcriptomes were sequenced. Differential gene expression analysis was performed between treatments (control vs. treated) for all tissues together and separately. Beforehand, a draft genome of *E. cicutarium* was assembled to use it as reference for the transcriptome analysis, and its DNA methyltransferase genes were characterized. We found that 5-azaC up-regulated chromomethylase *CMT1* across all treated samples, and the domain rearranged DNA methyltransferase *DRM2* in juvenile roots. Furthermore, adult leaves showed more differentially expressed genes between control and 5-azaC treated samples compared to juvenile leaves, supporting long term transcriptomic effects of a short exposure to 5-azaC at seed germination. At adult stage, leaves of individuals treated with 5-azaC exhibited up-regulation of genes involved in seed dormancy release, photoinhibition, and osmotic stress. Finally, gene co-expression network analysis revealed a module of co-expressed genes with differential gene expression linked to 5-azaC treatment in juvenile roots, that was enriched with genes involved in retrotransposon activity and in anthocyanin metabolism. Altogether, this study illustrates how the experimental treatment with 5-azaC at seed stage generates tissue- and age-specific transcriptional shifts, directly affecting gene regulation and potentially broadening phenotype variation in this fast-growing annual plant.

## Introduction

Epigenetic regulation comprises various interrelated mechanisms that can contribute to phenotypic variation without modifying the DNA sequence (Henderson and Jacobsen 2007), including histone and chromatin modification (Fransz and De Jong 2002, Du, Johnson et al. 2015), gene expression regulation mediated by small interfering RNA (siRNA) (Matzke and Mosher 2014), and genomic DNA methylation (Zilberman, Gehring et al. 2007). Regarding the latter, plants in particular show an extended DNA methylation machinery compared to other organisms: not only cytosines in CpG positions are sensitive to methylation, but also those in CHG and CHH positions (H being A, C, T nucleotides) (Law and Jacobsen 2010). In addition to the DNA methyltransferases *DNMTs*, which maintain DNA methylation in CpG positions and are evolutionarily conserved across eukaryotes, plant-specific chromomethylases (*CMTs*), and Domains Rearranged DNA Methyltransferases (*DRMs*) in cross-talk with histone marks modification, are in charge of *de novo* methylation in CHH and its maintenance in CHG positions as well (Cao and Jacobsen 2002; Bewick et al. 2017). Moreover, *de novo* cytosine methylation in any sequence context is carried out by *DRM2* through a siRNA-directed DNA methyltransferase pathway (RdDM) unique to plants (Cao, Aufsatz et al. 2003, Henderson, Deleris et al. 2010, Matzke and Mosher 2014). Finally, plants possess a unique mechanism of active DNA demethylation that involves a family of 5-methylcytosine specific DNA glycosylases, including DEMETER and Repressor Of Silencing (ROS), which play a critical role in modulating the activity of transposable elements (TEs) and gene expression (Law and Jacobsen 2010).

While DNA methylation in CpG can be a stable epigenetic mark in certain genomic regions (e.g. gene body methylation)(Bewick et al. 2017), being suitable even for the reconstruction and dating of shallow phylogenies (Yao et al. 2023), more dynamic changes can be observed in other genomic regions (Yao et al. 2023) and sequence contexts that routinely differed between tissues, organs and developmental stages (Widman et al. 2014; Urich et al. 2016; Bhatia et al. 2018; Gutzat et al. 2020; N’Diaye et al. 2020) or may change in response to stress (Gallusci et al. 2023 and references therein; Yao et al. 2023). Distinguishing between these two conditions makes the interpretation of experiments designed to understand the contribution of epigenetic regulation in plant adaptation very challenging in non-model plants, which usually lack good quality genomic resources (Richards et al. 2017). DNA methylation inhibitors have been used for decades to generate phenotypic innovations and investigate the association between DNA methylation and phenotypic plasticity (Dvořák Tomaštíková and Pecinka 2024). In particular, the cytidine analog 5-azacytidine (5-azaC) has been used to experimentally manipulate DNA methylation, providing the opportunity to decouple epigenetic regulation from other experimental factors suspected to be dependent on DNA methylation (Lang, Wang et al. 2017, Huang, Liu et al. 2019, Jia, Zhang et al. 2020, Osorio-Montalvo, De-la-Peña et al. 2020). As a cytidine analogue, 5-azaC is incorporated into the genomic DNA during each replication event, binding covalently to DNA methyltransferases (DNMTs), and therefore causing a decline in DNA methylation levels (Christman 2002, Stresemann & Lyko 2008). Several plant transcriptomic studies show how these genes are up-regulated by 5-azaC (Xu, Wang et al. 2017, Kiselev, Ogneva et al. 2019), whereas others report down-regulation (Zheng, Zhang et al. 2023). Unfortunately, the transcriptomic response to 5-azaC has been always studied in seedlings of model plants with small genomes (Griffin, Niederhuth et al. 2016, Kiselev, Ogneva et al. 2019, Xu, Wang et al. 2017, Zhang, Si et al. 2020), leaving wild non-model plant species in the dark. Still, 5’-azacytidine has been proven to reduce global DNA methylation levels in several plant species (Fieldes, Schaeffer et al. 2005, Browne, Mead et al. 2020, Osorio-Montalvo, De-la-Peña et al. 2020, Troyee, Medrano et al. 2022, Alonso, Medrano et al. 2025), as well as produce changes in phenotype (Prakash and Kumar 1997, Fieldes and Amyot 2000, Bossdorf, Arcuri et al. 2010, Munsamy, Rutherford et al. 2013, Alonso, Medrano et al. 2025). Thus, understanding the consequences of 5-azaC treatment on gene expression and developmental regulation in species with different genome features could be helpful to better understand its mode of action, potential side effects (see Dvořák Tomaštíková and Pecinka 2024 for a recent review), and the utility of 5-azaC exposure experiments for investigating the role of epigenetic regulation in plant adaptation to heterogeneous and stressful environments (Richards et al. 2017).

In this study, we exposed seeds of a single inbred line of the widespread Mediterranean herb *Erodium cicutarium* (Geraniaceae) to 5-azaC, following the same protocol that was known to reduce global DNA methylation in leaves and roots at adult stage (Alonso, Medrano et al. 2025). We extracted and sequenced RNA from juvenile roots, juvenile leaves and adult leaves, and compared the transcriptomes of treated and control plants. We hypothesized that (i) 5-azaC will affect the transcriptome in the three tissues analyzed, (ii) the effects will be stronger at the juvenile stage and (iii) the response will be likely more variable across individuals at the reproductive stage. In order to improve functional characterization of recorded changes, we also *de novo* assembled the genome sequence of *E. cicutarium* paying particular attention to DNA methyltransferase (DNMT) genes.

## Methods

### Study species and genome assembly

*Erodium cicutarium* (L.) L’Hér. is a widespread, herbaceous annual plant with a fast life-cycle and a high ability for autonomous selfing, that is frequently associated with disturbed and ruderal habitats (Fiz-Palacios et al. 2006). Its genome size has been estimated to range from 1.2 to 2.7 Gbps (Zonneveld 2019), different ploidy levels have been reported from different geographic regions, ranging from diploid to hexaploid (Fiz-Palacios et al. 2006, Zonneveld, 2019). The study material was collected in Cazorla mountains (Jaén province, SE Spain) where preliminary data suggest specimens have 2n = 40 chromosomes.

DNA was isolated from leaves of a single *E. cicutarium* individual, the F4 generation of a self-pollinated lineage (strain *F0-CH34_F3-Ecic127_F4-01*). Library preparation and sequencing were performed by AllGenetics (www.allgenetics.eu). Illumina paired-end short reads library were constructed using the HiSeq X Reagent Kit v2.5, sequenced in a HiSeq X PE150 (Illumina) with an Illumina patterned flow cell. PacBio long reads library was constructed using SMRTbell Express Template Prep Kit 2.0 (PacBio), sequenced with a Sequel II sequencer (PacBio), with an SMRT Cell 8M under the Long-reads mode. The *de novo* genome assembly of *E. cicutarium* is described thoroughly in Supplemental Information. In brief, short and long reads were assembled *de novo* into “mega-reads” using the software MaSuRCA v3.4.2 (Zimin, Marcais et al. 2013), then polished with POLCA (Zimin and Salzberg 2020). GenomeScope (Vurture, Sedlazeck et al. 2017) was used on the Illumina paired-end reads to infer genome length and ploidy. The quality and completeness of the genome assembly was evaluated using BUSCO V5.beta.1 (Simão, Waterhouse et al. 2015). Further information about quality check, gene prediction, and protein annotation is thoroughly explained in **Supplemental Information**.

### Characterization of DNA methyltransferases in E. cicutarium genome

Gene models annotated as DNA methyltransferases, chromomethylases, and domain rearranged DNA methyltransferases in the *E. cicutarium* genome assembly annotations were retrieved. Amino acid sequences encoding for DNA methyltransferases in *A. thaliana* genome were retrieved from UniProtKB (Consortium 2007), accessed on May 28, 2024 (accession numbers specified in **Table 2**). DNA methyltransferase protein sequences were also retrieved from the evolutionary close species *Geranium maculatum* (Geraniaceae) (Blanco & Leebens-Mack, unpub. data) using local BLASTp (Altschul, Gish et al. 1990, Camacho, Coulouris et al. 2009), which were curated with another BLASTp search against UniProtKB database (Consortium 2007). The amino acid sequences from *A. thaliana, G. maculatum*, and *E. cicutarium* were aligned with CLUSTAL Omega online toolset (Sievers, Wilm et al. 2011), and a neighbor-joining tree with 500 bootstrap permutations was built with MEGA v12.0.10 to infer the phylogenetic relationships of *E. cicutarium* DNA methyltransferases. A second alignment including exclusively *E. cicutarium DNMT, CMT*, and *DRM* protein sequences was used to produce sequence logos of the catalytic conserved motifs shown in (Pavlopoulou and Kossida 2007), using webLogo (Crooks, Hon et al. 2004).

**Table 1.**
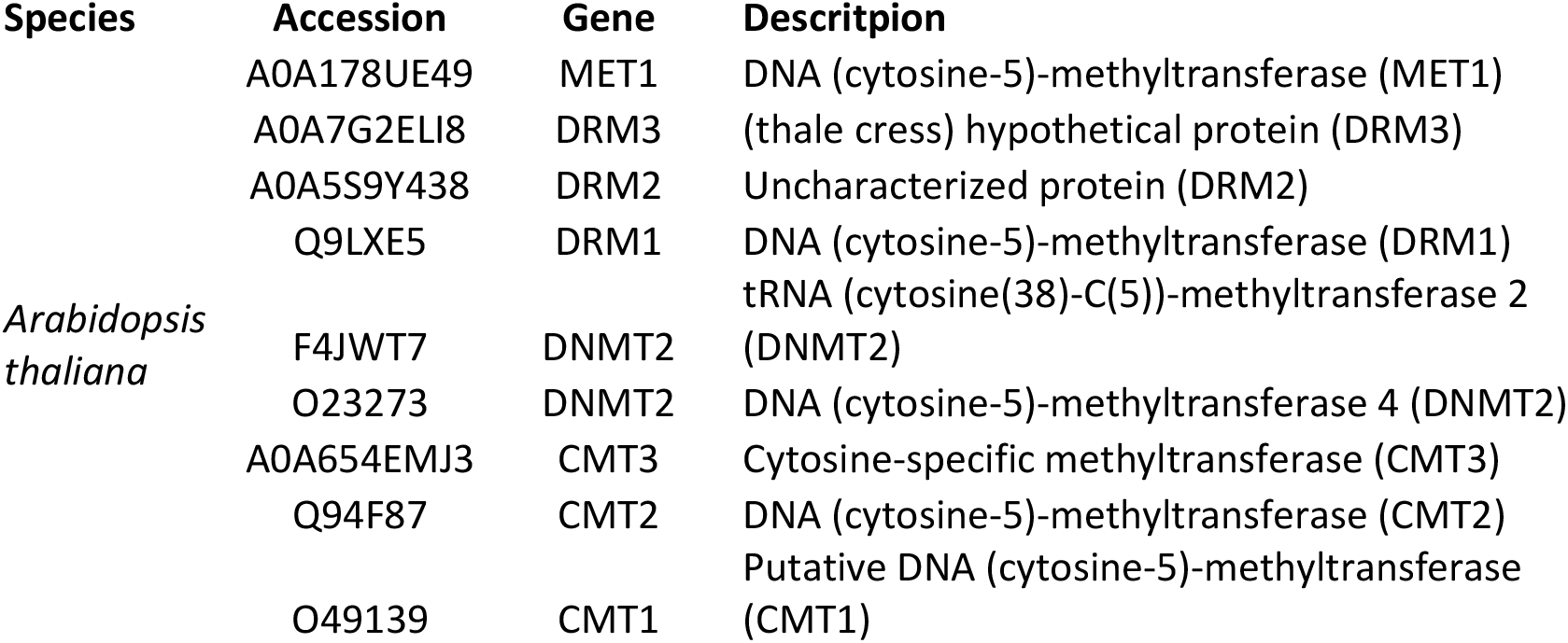
DNA methyltransferase amino acid sequences retrieved for phylogenetic analyses. Species name, UniProtKB accession number, gene name and description are shown in columns from first to fourth, respectively. Each rows shows a sequence entry.

**Table 2.** *Erodium cicutarium* RNA-seq library statistics. Columns show plant ID, sample ID (labeled after the individual), Sequencing Read Archive (SRA) accession number, seed treatment, tissue, developmental stage (age), million sequences in the raw library, million sequences after trimming, and the percentage of reads aligned to the *E. cicutarium* genome.

| Plant ID | Sample ID | Accession | Treatment | Tissue | Age | M seq (raw) | M seq (trim) | % Aligned |
| --- | --- | --- | --- | --- | --- | --- | --- | --- |
| ECI1 | ecic01_C_L_Y | SRR29751979 | Control | Leaf | Juvenile | 39.8 | 38.6 | 77.20 |
| ECI1 | ecic01_C_R_Y | SRR29751978 | Control | Root | Juvenile | 42 | 41.2 | 84.95 |
| ECI2 | ecic02_A_L_Y | SRR29751967 | Azacytidine | Leaf | Juvenile | 35.8 | 35 | 85.71 |
| ECI2 | ecic02_A_R_Y | SRR29751963 | Azacytidine | Root | Juvenile | 26.8 | 26.2 | 84.73 |
| ECI3 | ecic03_C_L_Y | SRR29751962 | Control | Leaf | Juvenile | 39.4 | 38.2 | 73.30 |
| ECI3 | ecic03_C_R_Y | SRR29751961 | Control | Root | Juvenile | 31.8 | 30.8 | 82.47 |
| ECI4 | ecic04_A_L_Y | SRR29751960 | Azacytidine | Leaf | Juvenile | 31.4 | 30.8 | 85.71 |
| ECI4 | ecic04_A_R_Y | SRR29751959 | Azacytidine | Root | Juvenile | 34.6 | 34 | 84.71 |
| ECI5 | ecic05_C_L_Y | SRR29751958 | Control | Leaf | Juvenile | 34.8 | 34.2 | 84.21 |
| ECI6 | ecic06_A_L_Y | SRR29751957 | Azacytidine | Leaf | Juvenile | 28 | 27.6 | 84.78 |
| ECI6 | ecic06_A_R_Y | SRR29751977 | Azacytidine | Root | Juvenile | 37.4 | 36.6 | 84.15 |
| ECI7 | ecic07_C_L_Y | SRR29751976 | Control | Leaf | Juvenile | 40.2 | 39.4 | 82.23 |
| ECI7 | ecic07_C_R_Y | SRR29751975 | Control | Root | Juvenile | 39 | 38.6 | 86.01 |
| ECI8 | ecic08_A_L_Y | SRR29751974 | Azacytidine | Leaf | Juvenile | 40.6 | 39.8 | 86.43 |
| ECI8 | ecic08_A_R_Y | SRR29751973 | Azacytidine | Root | Juvenile | 42 | 41.2 | 85.44 |
| ECI9 | ecic09_C_L_Y | SRR29751972 | Control | Leaf | Juvenile | 41.8 | 41.4 | 84.54 |
| ECI9 | ecic09_C_R_Y | SRR29751971 | Control | Root | Juvenile | 41.8 | 41 | 81.46 |
| ECI10 | ecic10_A_L_Y | SRR29751970 | Azacytidine | Leaf | Juvenile | 38 | 37 | 80.54 |
| ECI10 | ecic10_A_R_Y | SRR29751969 | Azacytidine | Root | Juvenile | 42.2 | 41.2 | 83.01 |
| ECI11 | ecic11_C_L_Y | SRR29751968 | Control | Leaf | Juvenile | 34 | 33 | 82.42 |
| ECI11 | ecic11_C_R_Y | SRR29751966 | Control | Root | Juvenile | 34.8 | 33.8 | 84.62 |
| ECI12 | ecic12_A_L_Y | SRR29751965 | Azacytidine | Leaf | Juvenile | 39 | 38 | 85.79 |
| ECI12 | ecic12_A_R_Y | SRR29751964 | Azacytidine | Root | Juvenile | 36.8 | 36 | 85.56 |
| E1 | ecic1_A_L_O | SRR29751797 | Azacytidine | Leaf | Adult | 42 | 41.6 | 82.21 |
| E12 | ecic12_C_L_O | SRR29751828 | Control | Leaf | Adult | 41.8 | 41.4 | 86.96 |
| E13 | ecic13_A_L_O | SRR29751827 | Azacytidine | Leaf | Adult | 41.8 | 41.4 | 86.96 |
| E14 | ecic14_C_L_O | SRR29751816 | Control | Leaf | Adult | 41.8 | 41.4 | 83.57 |
| E17 | ecic17_A_L_O | SRR29751805 | Azacytidine | Leaf | Adult | 42.2 | 41.8 | 83.73 |
| E18 | ecic18_C_L_O | SRR29751798 | Control | Leaf | Adult | 42 | 41.6 | 84.13 |
| E2 | ecic2_C_L_O | SRR29751793 | Control | Leaf | Adult | 42 | 41.6 | 83.65 |
| E21 | ecic21_A_L_O | SRR29751796 | Azacytidine | Leaf | Adult | 42 | 41.6 | 86.06 |
| E24 | ecic24_C_L_O | SRR29751795 | Control | Leaf | Adult | 42.2 | 41.8 | 84.69 |
| E29 | ecic29_A_L_O | SRR29751794 | Azacytidine | Leaf | Adult | 42 | 41.6 | 83.65 |
| E5 | ecic5_A_L_O | SRR29751813 | Azacytidine | Leaf | Adult | 41.8 | 41.4 | 83.57 |
| E8 | ecic8_C_L_O | SRR29751799 | Control | Leaf | Adult | 42.2 | 41.8 | 85.65 |

### Experimental exposure to 5-azaC

Twelve seeds coming from a single maternal *Erodium cicutarium* plant were scarified with sandpaper, then soaked in 200 µL of either a control or a 500 µM 5-azaC (Sigma A2385–100mg, initially diluted to 100 mM in Sigma DMSO) solution for two days (Alonso, Medrano et al. 2017). For the control solution, the same proportion of DMSO was diluted in water (3:97; v:v). Control and treated seeds were afterwards germinated individually in seedling nursery trays (4x4x7 cm) and grown in a climatic chamber (Aralab CLIMAPLUS 400; with 12 h light and 22°C : 18°C) for five weeks until they reached juvenile stage, i.e., plants that have the cotyledons and the first four true leaves. Another group of 12 plants from a different experimental set (with identical 5-azaC and control exposure) were left to grow up over 12 weeks in the greenhouse (10 h light; 25°C : 15°C) to reach an adult stage, i.e., plants that have starting to open their first flowers to assess whether some transcriptomic effects after seed treatment persist up to the stage that could be transmitted to gametes.

### RNA extraction, sequencing, quality check and alignment to genome

Fresh juvenile leaf (N = 12), juvenile root (N = 12) and adult leaf (N = 12) samples (10 to 100 mg) from each experimental plant were flash frozen in liquid nitrogen and stored at -80 °C. Both RNA extraction and sequencing were performed by Macrogen Inc. (Seoul, Republic of South Korea). RNA root sample from juvenile individual *ecic05* did not pass RNA isolation quality check, and was discarded from sequencing. Paired-end reads of 151 bp length RNA-seq libraries were generated using Truseq Stranded mRNA Library Prep kit (20020595, Illumina), and Illumina sequencing was performed using a NovaSeq6000 machine. Reads are available at NCBI sequence read archive (SRA), under the BioProject accession numbers PRJNA1128899 (juvenile samples) and PRJNA1129331 (adult samples).

Read quality check was assessed with FastQC (Andrews 2010) for each step of the pipeline. Reads were trimmed out of low quality positions and Illumina adapters with Trimmomatic v0.39 (Bolger, Lohse et al. 2014), using the options ‘*-phred33 ILLUMINACLIP:$adapter_fasta_file:2:30:10 LEADING:28 TRAILING:28 SLIDINGWINDOW:4:15 MINLEN:30*’, and quality checked again with FastQC. Trimmed reads were aligned to the draft genome of *E. cicutarium* with STAR v2.7.10 (Dobin, Davis et al. 2012), with options ‘*--sjdbOverhang 149 --quantMode TranscriptomeSAM GeneCounts --twopassMode Basic -- outSAMunmapped Within --outSAMtype BAM SortedByCoordinate*’, and again quality checked with FastQC. FastQC files were scanned with MultiQC v1.13.dev0 (Ewels, Magnusson et al. 2016) to summarize the number of reads and calculate fragment alignment rate to genome. Counts of fragments (paired reads) mapped to gene models were extracted with *featureCounts* v2.0.3 command from the Subread package (Liao, Smyth et al. 2013), with options ‘*-p -s 0 -g Parent -t exon*’.

### Differential gene expression analysis

Genes in *E. cicutarium* genome with less than 100 fragment counts across all samples were discarded from subsequent analyses. Remaining raw fragment count per gene values were normalized using the DESeq2 median of ratios method (Love, Huber et al. 2014), and differential gene expression analysis was performed using DESeq2 by fitting expression data to the formula *expression ∼ 0 + group*, (*group* consisted in six levels: control leaves from juvenile, control leaves from adult plants, 5-azaC leaves from juvenile, 5-azaC leaves from adult plants, control roots from juvenile, and 5-azaC roots from juvenile). Differential expression analysis was performed to test the effect of 5-azaC. We compared all controls samples with all 5-azaC samples, followed by separate within tissue and within age comparisons between control and 5-azaC samples. A gene was considered a differentially expressed gene (DEG) when its false discovery rate (FDR (Benjamini and Hochberg 1995)) in any of the comparisons was below 0.05. The DEGs related to epigenetic regulation processes (including DNA methylation, chromatin modification, and siRNA regulation) are commented in the results. Overlapping DEGs between different DEG sets were tested to check whether the overlap was greater than expected by chance with the hypergeometric test using the *phyper* R function. All statistical analyses were performed in R v4.3.1 (R Core Team 2023).

### Gene co-expression network analysis

In addition to gene expression analysis, weighted gene co-expression networks were analyzed with the R package *WGCNA* (Langfelder and Horvath 2008) to generate groups of co-expressed genes (modules) associated to the control vs. 5-azaC treatment. Gene modules were determined using the hybrid method implemented in the *blockwiseModules()* function, using the parameters ‘*maxBlockSize = 70973, networkType = “unsigned”, power = 6, detectCutHeight = 0*.*9, deepSplit = 4, minModuleSize = 50*’. To evaluate connectivity of genes within their modules, we calculated Kleinberg’s hub centrality scores for all genes with the *hub_score()* function from *igraph* R package (Csárdi and Nepusz 2006). PCA of module eigengenes (MEs) was performed to explore sample clustering and their relationships with the modules. To test the association between modules and treatments, module gene significances were calculated by transforming each module gene’s FDR as -log_10_(FDR) and calculating its mean per module. Given there are four comparisons between control vs. 5-azaC treatment (all samples, juvenile roots, juvenile leaves, and adult leaves), each module showed four module gene significance values. Gene module memberships from modules with the highest gene significances were calculated as the resulting Pearson’s correlation coefficient from correlations between gene expression and its relevant module eigengenes (see **Supplemental Information** for exceptions).

### Gene ontology term enrichment analysis

Gene ontology (GO) terms associated to DEGs and WGCNA modules were retrieved from the *E. cicutarium* draft genome annotation files. GO term enrichment analysis was performed with *topGO* (Alexa and Rahnenfuhrer 2023) R package by comparing the GO terms from all expressed genes with those from the DEG sets, using the weighted Fisher’s exact test (FET) statistic. A GO term was considered enriched when the FDR adjusted value of the weighted FET was less than 0.05. When enriched GO term sets were large enough (hundreds), GO terms and their FDR values were analyzed with REVIGO to improve visualization (Supek, Bošnjak et al. 2011), using 0.5 set and UniProtKB as reference protein database.

## Results

### Erodium cicutarium draft genome features

Illumina sequencing yielded 100 Gbp from 333.20 Million paired-end 150 bp reads, PacBio sequencing yielded a total of 131.55 Gbp from 6.75 million long reads. Both Illumina and PacBio reads can be found in the NCBI’s SRA, under BioProject accession PRJNA984161. The assembly was built with an estimated sequencing coverage of 163.68x. GenomeScope inferred a genome sequence length of 765.02 Mbp (**Figure 1A**), and produced a profile compatible with diploid and allotetraploid genomes (Ranallo-Benavidez, Jaron et al. 2020)). The reference genome assembly for *E. cicutarium* ended up with a total length of 801.44 Mbp (∼5 % longer than predicted by GenomeScope), distributed in 1,206 contigs, with a contig maximum length of 9.13 Mbp and a N50 of 1.95 Mbp. Genome assembly’s average GC content was 41 %, and the repeated elements (REs) content in the assembly was 41.97 %. Most contigs and scaffolds show a high frequency of genic regions, showing evenly distributed RNA-seq coverage along the contigs that, together with a lower frequency of repetitive elements. The raw reads are hosted in the SRA from NCBI, under accession number SRR28842229. The genome sequence is hosted in the genome repository of NCBI, under accession number JBDFRG000000000.

**Figure 1.**
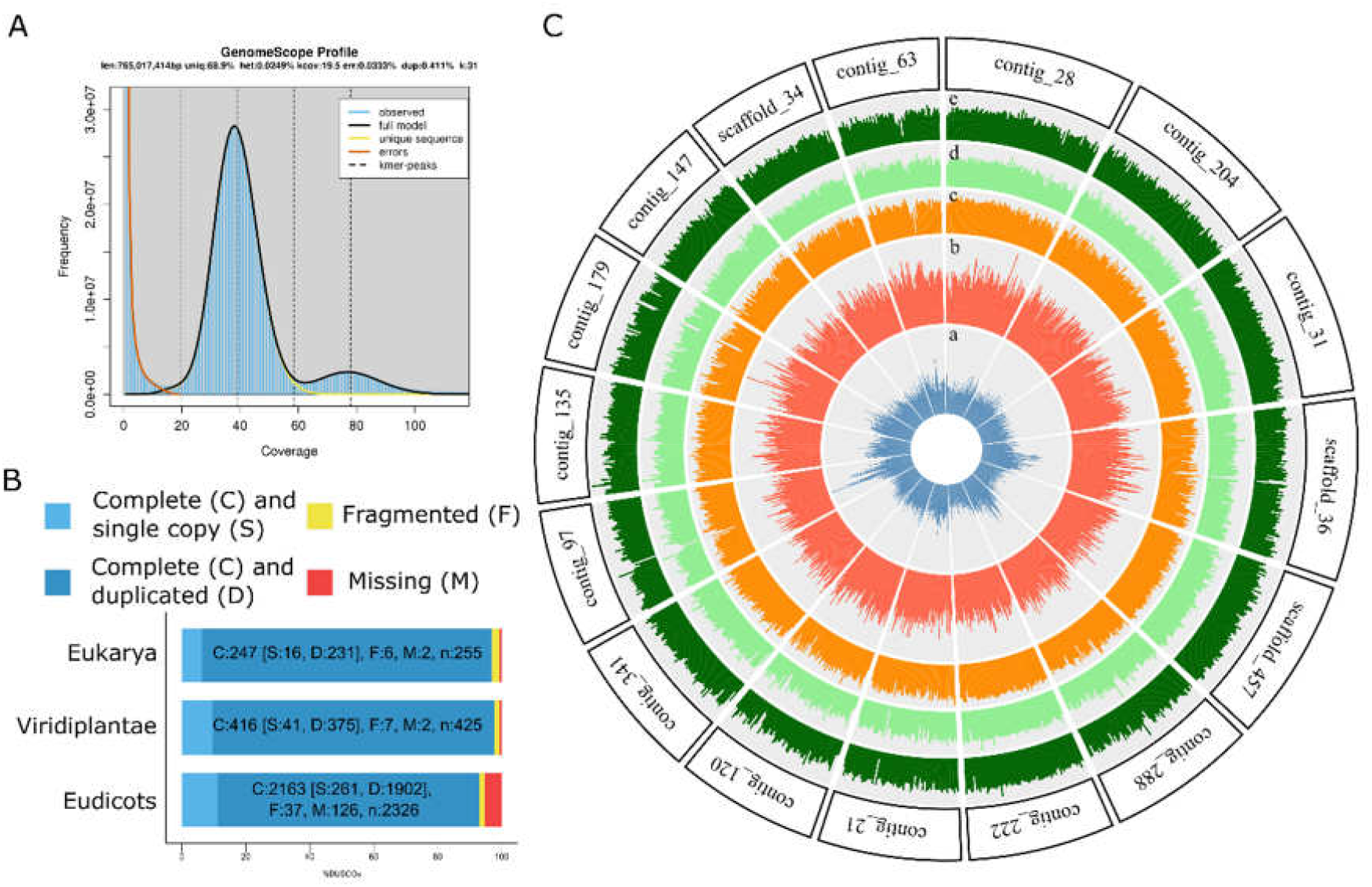
*E. cicutarium* genome assembly quality check. A: GenomeScope plot from filtered Illumina reads for k-mer 31. X-axis shows coverage values, Y-axis shows coverage distribution frequency for the observed (blue bars) and expected (black line) genome models. Estimated coverage by Illumina reads was 130x. The two peaks, the first being higher than the second, indicate polyploidy. The header shows the estimated genome length (len), percentage of unique k-mers (uniq), estimated percentage of heterozygosity (het), mean k-mer coverage (kcov), sequencing error rate (err), and length of the analyzed k-mers (k). B: BUSCO completeness analysis of the genome. X-axis shows percentage values, Y-axis shows BUSCOs found in the Eukarya, Viridiplantae and Eudicots databases, colored bars show the number of BUSCOs found in each search: single copy complete (light blue), duplicated complete (dark blue), fragmented (yellow) and missing (red). C: Circos graph of the genome. The outer ring shows the contigs with a length of 6 Mbps or more (n=16), inner rings show repeated element (a) and gene (b) density in blue and red, respectively; and RNA-seq fragment coverage of juvenile roots (c), juvenile leaves (d), and adult leaves (e) in orange, light green and dark green, respectively.

The total number of inferred gene models was 186,428, with 172,514 showing an open reading frame. Genome completion assessed through BUSCO score casted a 92.99 % of complete sequences against the *eudicot_obd10* database (accessed on February 2, 2023, Figure 1B), with a proportion of duplicated complete BUSCOS of 87.93 %; a similar trend was found when comparing against Viridiplantae and Eukarya databases. Figure 1C shows contigs longer than 6 Mbp with repeated elements and gene densities, and RNA-seq expression of the samples.

### DNA methyltransferase sequence analysis

Genome annotation predicted a total of 22 DNA methyltransferase genes: nine chromomethylases (*CMT*), four DNA methyltransferases (*DNMTs*), and nine domain rearranged DNA methyltransferases (*DRMs*). BLASTp search retrieved 15 homologous sequences (five *CMTs*, five *DNMTs*, and five *DRMs*) from *G. maculatum* genome’s protein set. Although no *MET1* homologous sequence was found in *E. cicutarium* nor *G. maculatum* genomes, analysis of both Geraniaceae and *A. thaliana* DNA methyltransferase sequences revealed that a clade including four *E. cicutarium* sequences annotated as *DNMTs* and five *G. maculatum DNMT* sequences were as closely related to *A. thaliana MET1* as to the *DNMT2* with accession AT4G14140 (**Figure 2**). The phylogeny also shows a loss of the *CMT1* clade and a diversification of clade CMT3 in both Geraniaceae. Alignment of amino acid sequences also confirmed the presence of the six conserved catalytic domains described so far for plants (Pavlopoulou and Kossida 2007) in the *E. cicutarium* DNA methyltransferase gene set **Supplemental Figure S1**. The amino acid sequences used for the alignment are available in **Supplemental Information**.

**Figure 2.**
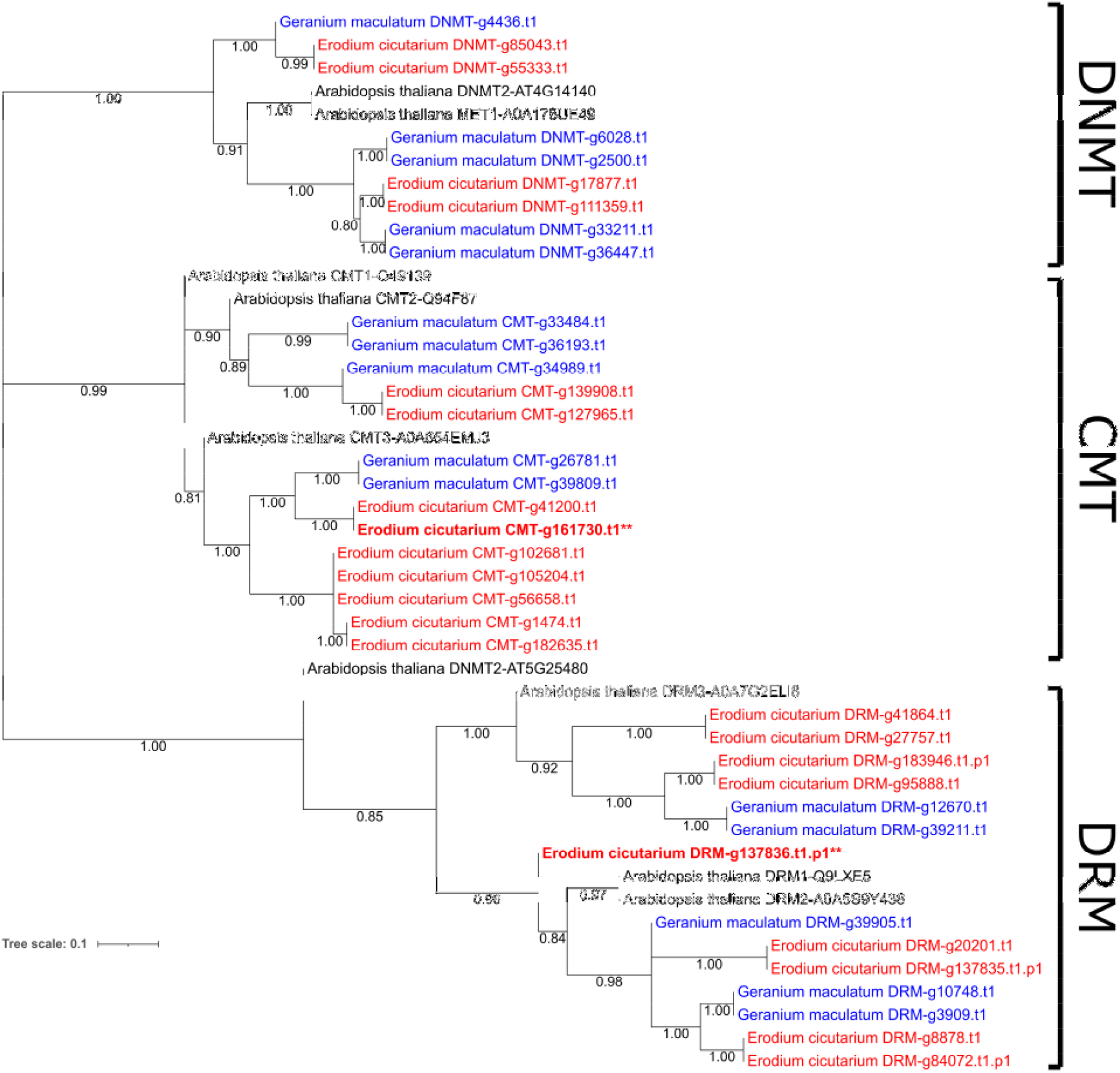
Characterization of DNA methyltransferases from *E. cicutarium* genome. The figure shows the consensus Neighbor-joining tree generated by MEGA v12.0.10 from an alignment between the amino acid sequences of the 22 DNA methyltransferases annotated in the *E. cicutarium* (red) draft genome, 15 from the *G. maculatum* genome protein set (blue), and those retrieved from UniProtKB for *A. thaliana* (black) DNA methyltransferases. The scale bar denotes a length of 0.1 substitutions per nucleotide site. Bootstrap branch supports are shown in each node. The main three DNA methyltransferase clades (domains rearranged methyltransferases *DRM*, DNA methyltransferases *DNMT*, and chromomethylases *CMT*) are marked with delimitation bars in the right part of the tree. Sequences whose gene was found differentially expressed in any control vs. 5-azaC treatment comparison is marked in bold, asterisks indicate the statistical significance of such difference (^**^: 0.01 > FDR > 0.001).

### RNA sequencing and mapping to genome

A total of 1,356 million pair-end reads of 151 bps length were produced by the Illumina RNA sequencing experiment (N = 35 samples; **Table 3**). After trimming, 1,332.6 million pair-end reads remained, with lengths ranging from 32 to 151 bps. Mean read mapping rate to *E. cicutarium* draft genome was 83.8 %, ranging from 73.3 % to 87.0 %. After fragment count summarizing and expression value filtering, 70,973 gene models were considered for differential gene expression analysis. **Table 3** shows all the sequencing statistics per library.

### Differential gene expression analysis

DESeq2 normalization managed to standardize the median of the fragment counts **(Supplemental Figure S2A)**. Principal component analysis (PCA) of the top 500 most expressed genes across samples predicted strong effects of tissue and age in gene expression, and not so obvious divergence for the comparison between control and 5-azaC treated plants that was the focus of our experiment **(Supplemental Figure S2B)**. As expected, there were tens of thousands of DEGs between juvenile roots and juvenile leaves (54,424), and juvenile and adult leaves (37,102). Below we describe divergence associated to 5-azaC seed treatment **(Supplemental Figure S3, Supplemental Table S1)**, while gene expression divergence between either leaves and roots or juvenile and adult leaves is presented in **Supplemental Information (Supplemental Figure S3)**.

DESeq2 detected 164 DEGs for the control and 5-azaC comparison (147 up-regulated and 17 down-regulated by 5-azaC). Among the DEGs, a chromomethylase *CMT1* homologous gene (involved in non-CpG methylation maintenance), and three *MYB* transcription factors (involved in stomatal development, jasmonate and cytokinins signaling pathways, and other processes) were upregulated by 5-azaC, as well as some genes related to anthocyanin synthesis pathway (*B-MYB*, UDP-glycosyltransferase 84B1). Hypergeometric test showed that the overlap of the DEG sets global control vs 5-azaC was not greater than expected by chance in any case (P-value > 0.05)

Differential gene expression analyses of 5-azaC effect within tissue datasets detected 99 and 487 DEGs for juvenile leaves and juvenile roots, respectively **(Figure 3, Supplemental Tables S2 and S3)**. In juvenile leaves, 59 DEGs were up-regulated in the 5-azaC treated samples, and 40 down-regulated (including a *histone H3*.*2* protein, involved in chromatin modeling). In juvenile roots, 464 DEGs were up-regulated in the 5-azaC treated samples (including a domain rearranged methyltransferase *DRM2* homolog, involved in non-CpG methylation maintenance; and a DNA-directed RNA polymerase 23kD subunit, involved in siRNA synthesis and subsequent RNA-directed DNA methylation), and 23 down-regulated. Comparison within adult leaves detected 187 DEGs, 142 up-regulated and 45 down-regulated by 5-azaC **(Figure 3, Supplemental Table S4)**. The number of overlapping DEGs between these datasets still was not greater than expected by chance in any case (P-value > 0.05), suggesting rather distinct effects of 5-azac in the three sample-types here analyzed.

**Figure 3.**
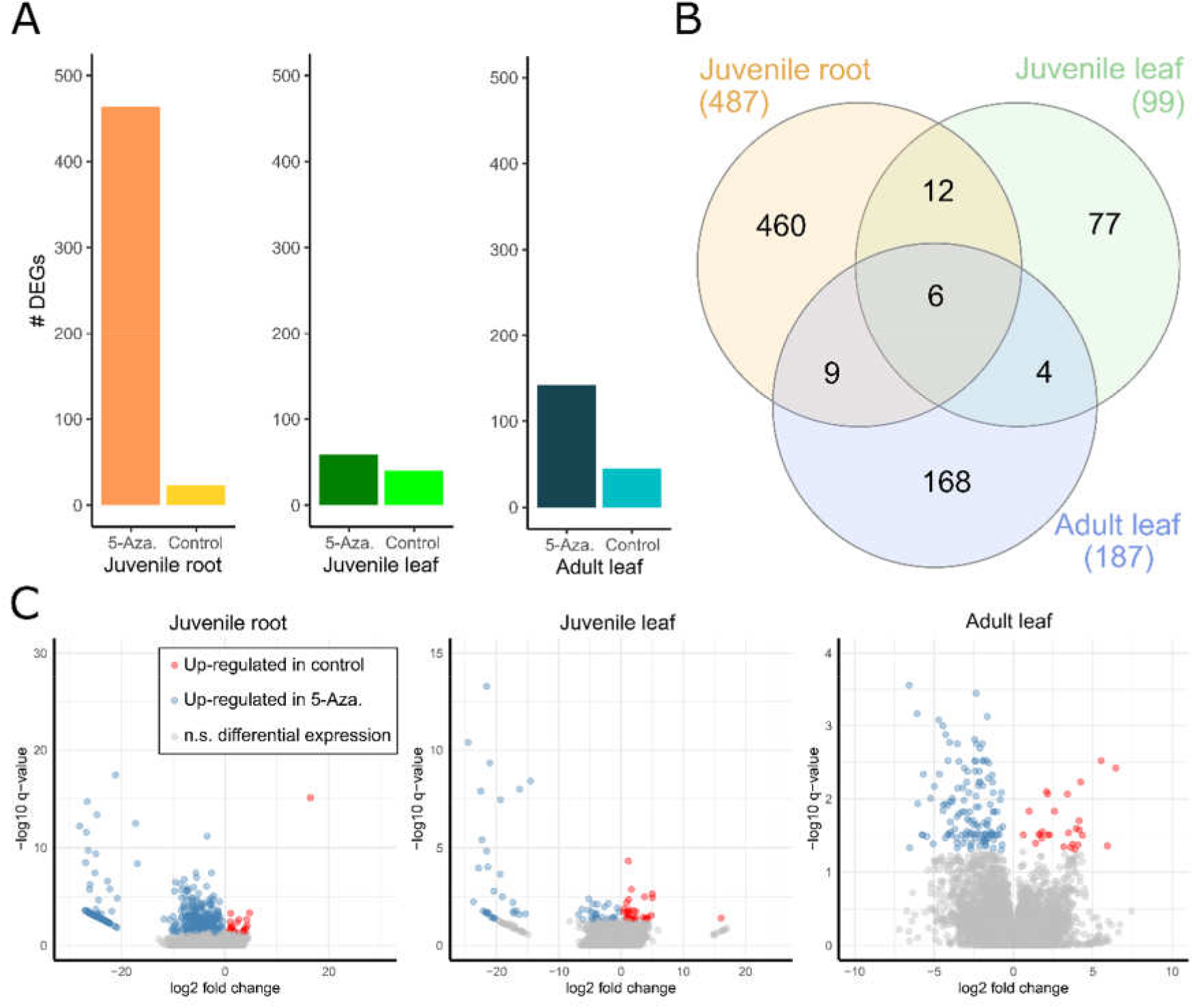
Differential gene expression analysis of 5-azacytidine treatment within tissue and age samples. A: number of differentially expressed genes (DEGs) between control and 5-azacytidine within young leaves, young roots and adult leaves. X-axis shows all tested pairs of groups, Y-axis show the number of DEGs up-regulated in each group. B: Venn diagram showing the number of overlapping DEGs per group comparison. Each circle represents a comparison between two groups, numbers inside circles show how many DEGs are in each overlapping subset. Numbers in parentheses show the total number of DEGs found in the comparison. C: Volcano plots of each tested comparison (specified in the header of each plot). X-axis shows the logarithm to the base 2 of the fold change, Y-axis shows the negative logarithm to the base 10 of the false discovery rate, blue and red dots represent genes down-regulated (blue) or up-regulated (red) by 5-azacytidine in each comparison, while grey dots are genes not differentially expressed in the comparison.

**Supplemental Figure S4** shows the gene expression profiles of representative genes of the two clades of *DNMT* (phylogenetically closest sequences to *A. thaliana MET1* and *DNMT2* in *E. cicutarium* genome) that were both particularly more expressed in roots, as well as the two differentially expressed *CMT1* and *DRM2* gene copies from *E. cicutarium* genome. Note that despite not showing differential expression between control and 5-azaC treated samples, the two *DNMT2* genes showed a higher variation in expression across samples treated with 5-azaC compared to controls. Despite *CMT1* being mainly expressed in roots, it was up-regulated across all 5-azaC treated samples (FDR = 0.004), due to the control samples having more *CMT1* expression values equals to zero in the two leaf-types. Finally, *DRM2* was up-regulated in 5-azaC treated roots (FDR = 0.006), and despite showing higher expression values in juvenile leaves treated with 5-azaC compared to controls, the test was not statistically significant (FDR = 0.373).

### Gene ontology term enrichment analysis

A total of 16,072 GO terms were associated with the ca. 71,000 gene models included in the differential gene expression analysis. The DEG set from the 5-azaC general effect comparison did not show significant enriched GO terms, while DEG sets from the within tissue and age comparisons showed more variable outputs, with an overall trend for the sets from up-regulated genes in 5-azaC treated samples. In particular, in the control vs. 5-azaC comparison for juvenile leaves, no GO terms were enriched. In juvenile roots, DNA integration (GO:0015074) and another four GO terms linked to retrotransposon activity were enriched in the DEG set from roots treated with 5-azaC **Supplemental Figure S5A, Supplemental Table S5**. In adult leaves, the DEG set from samples treated with 5-azaC showed nine enriched GO terms, including two related with release from seed dormancy (GO:0048838, GO:1902039), positive regulation of gibberellic acid mediated signaling pathway (GO:0009939), response to osmotic stress (GO:0006970) and other functions associated to membrane transport and photo inhibition **(Supplemental Figure S5B, Supplemental Table S5)**. DEG sets from control up-regulated in root and adult leaf samples did not show enriched GO terms.

### Gene co-expression network analysis

A total of 57,411 genes were included in one of the twelve generated gene co-expressed modules **Supplemental Figure S6A, Supplemental Table S6**. PCA of module eigengenes (MEs, **Supplemental Table S7**) showed a similar clustering than the PCA with expression data, samples being separated by age and tissue rather than discriminating those differing in 5-azaC treatment **Supplemental Figure S6B**. Module 10 was excluded from this analysis due to excess of genes with a fragment count of zero (see **Supplemental Information** for the details on this decision). Thus, module M6 showed the highest gene significance values when considering control vs. 5-azaC adjusted p-values (**Figure 4**), being particularly associated with increased expression in samples treated with 5-azaC. It was enriched with GO terms related to retrotransposon activity and anthocyanin metabolism **Figure 4, Supplemental Table S8**. For the most specific control vs 5-azaC comparisons conducted for each sample-type, we found that the highest gene significance values for the juvenile roots was also held by module M6, whereas juvenile leaves and adult leaves comparisons were most associated to modules M9 and M12, respectively **Supplemental Figure S7**. Finally, the two differentially expressed DNA methyltransferase genes, *CMT1* and *DRM2*, were included in modules M1 and M2, respectively; both modules showed MEs up-regulated in roots, independently of their 5-azaC treatment.

**Figure 4.**
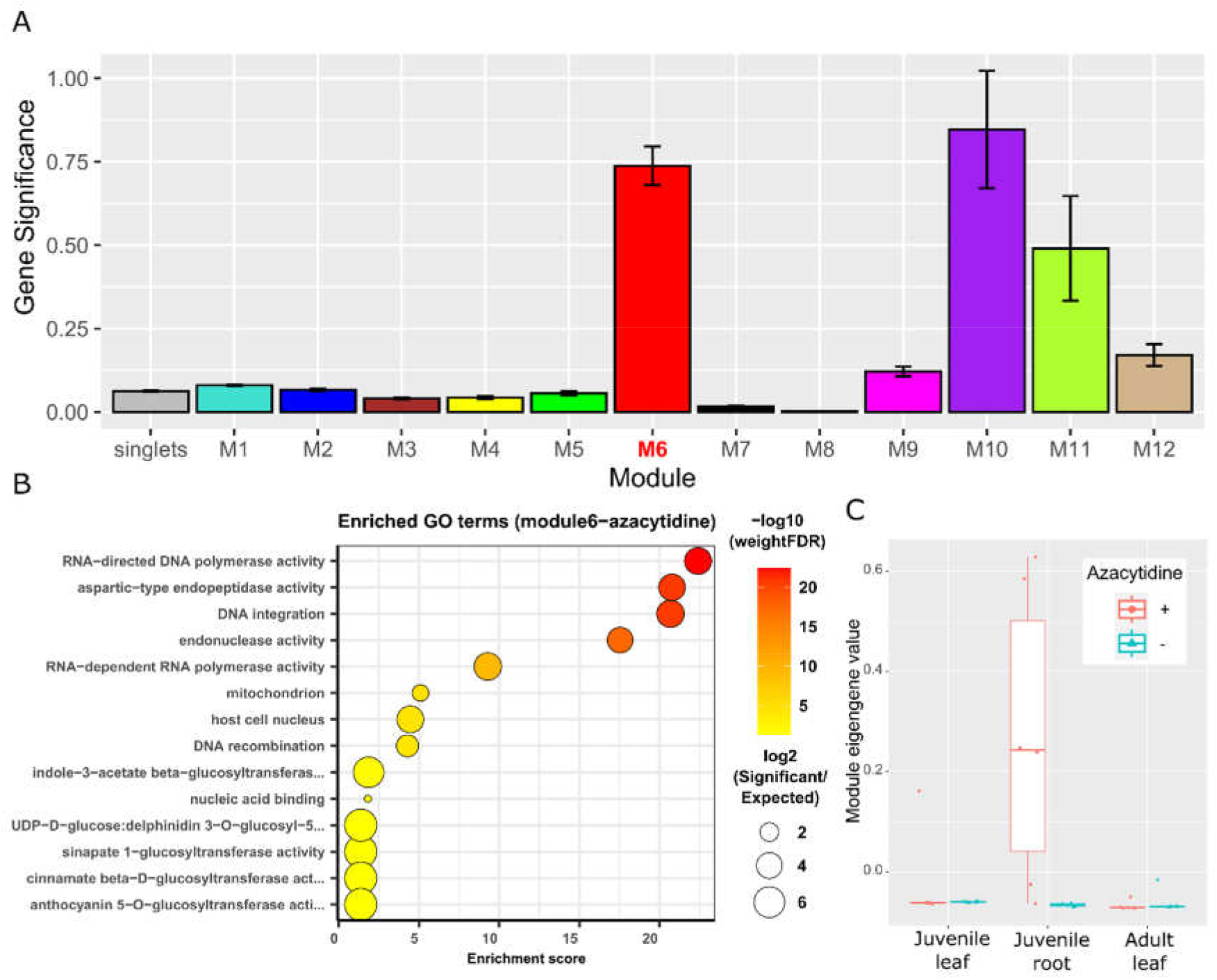
Gene co-expression network analysis and 5-azacytidine associated module M6 description. A: gene significances by module. X-axis shows the module name, Y-axis shows the gene significance (negative logarithm to the base 10 of the false discovery rate) values. Bars show the mean gene significance value per module, with error bars showing the standard errors. B: Enriched GO terms in genes from module M6. X-axis shows the enrichment score calculated by *topGO*, Y-axis shows the names of the enriched GO terms. Circles show the logarithm to the base two of the ratio between the observed and expected GO term counts (circle size) and the negative logarithm to the base 10 of the false discovery rate value (circle color). C: Eigengene values from module M6. X-axis shows each sample type, Y-axis shows the MEs. Each box shows percentiles 25, 50 and 75 of its respective group (with 5-azacytidine treated samples in red and control samples in blue).

## Discussion

Cytidine analogs such as 5-azaC have been established as relevant experimental tools for understanding DNA demethylation in model and non-model plant species (see Dvořák Tomaštíková and Pecinka 2024 for a recent review). Combined with another environmental factor of interest including biotic interactions or abiotic conditions, they provide a tractable strategy to detect direct links between epigenetic regulation and functional phenotypic responses in plant species with contrasting life histories or ecological features, contributing, thus, to the rising interest on ecological epigenetics (Alonso, Ramos-Cruz et al. 2019, Balao, Paun et al. 2018, Williams, Miller et al. 2023). Thus, understanding how 5-azaC affects gene expression and gene regulatory networks in different plant species is key for an accurate interpretation of its multiple phenotypic effects and potential contribution to uncover the epigenetic contribution to plant adaptation in heterogeneous and stressful environments. In the following paragraphs, we will discuss the transcription effects recorded in this study aimed to understand developmental and tissue-specific changes in the response to seed demethylation in *E. cicutarium*.

Exposure to 5-azaC can be toxic for plants (Qiu, Hother et al. 2010, Grzybkowska, Morończyk et al. 2018, Osorio-Montalvo, De-la-Peña et al. 2020), thus, adjusting the treatment concentration is critical to prevent increased mortality or dwarfism (Alonso et al. 2017 and references therein). Our study uses a 500 µM concentration, which is in range of other studies that did not intend to generate toxic effects on treated plants but were proved to significantly reduce DNA methylation (Troyee et al. 2022, Alonso et al. 2025). In fact, gene expression differences related to tissue and development stage overwhelm those related to 5-azaC, which increased individual variation in gene expression, rather than affecting it in a directional or unimodal way. Signs of stress response in the transcriptome did show only in adult leaves treated with 5-azaC, where GO terms from photoinhibition and osmotic stress were enriched. This trend may be a direct consequence of the 5-azaC toxicity or an indirect effect due to trade-offs induced by phenotype changes, but since the juvenile stages did not show such stress response in this study, the later option seems the more plausible.

Previous studies showed that 5-azaC treatment at seed stage successfully dropped global DNA methylation in *E. cicutarium* adult roots, but did affect more strongly to juvenile than adult leaves (Alonso, Medrano et al. 2025). Still, many DNA methylation changes were observed in adult leaves, which recorded a higher reduction of methylation in CG sequences and similar numbers of methylation gains and losses in cytosines at other sequence contexts (Balao et al. 2024).

5-azaC is expected to affect DNA methyltransferase gene expression, and several plant studies show how these genes are up-regulated by 5-azaC. In 8-week old leaves of *A. thaliana*, resulting hypomethylation from exposure to 5-azaC induced the up-regulation of DNA methyltransferases *MET1* (a specific *A. thaliana DNMT*), *CMT3, DRM1* and *DRM2*, together with other genes involved in the DNA repair pathway (Kiselev, Ogneva et al. 2019). Calluses of the *Citrus paradisi*, when exposed to 5-azaC, showed increased levels of *MET1* and *CMT1* transcripts (Xu, Wang et al. 2017). On the other hand, DNA methyltransferases *CMT3, MET1* and *DRM2* were down-regulated when exposed to 5-azaC in *Paeonia suffruticosa* (Zhang, Si et al. 2020). In our study, we did not find up-regulation of *DNMT* genes induced by 5-azaC, but found *CMT1* up-regulated instead, which might point towards alternative compensation mechanisms when DNA methylation in CG positions is inhibited. This lack of expression differences in *DNMTs* between treatments may indicate that the *DNMT* transcripts are consistently expressed whether the *DNMT* proteins are functional or inhibited, involving a lack of a regulatory mechanism to inhibit *DNMT* gene transcription under 5-azaC treatment. In addition, *DRM2* (involved in *de novo* cytosine methylation through the small RNA dependent DNA methylation pathway) and a *RdDM* polymerase component were also up-regulated in roots treated with 5-azaC, suggesting that plant specific *RdDM* pathway may be enhanced after 5-azaC treatment. *CMT1* and *DRM2* promote and maintain non-CpG methylation (Cao, Aufsatz et al. 2003), and are more sensitive to environmental cues (Du, Zhong et al. 2012, Bouyer, Kramdi et al. 2017), which may allow the treated plants some environmental-dependent responsivity for DNA methylation regulation, and potentially compensate for *DNMT* inhibition.

Juvenile roots were particularly susceptible regarding their transcriptome response to 5-azaC exposure: they did not only show more up-regulated DEGs by 5-azaC compared to juvenile and adult leaves, they also showed a strong association with module M6 of the gene co-expression network. Both DEGs in roots treated with 5-azaC and genes in module M6 contained gene sets enriched with transposable elements. It is well established that 5-azaC treatment up-regulates transposon expression in other plants (Hudson, Luo et al. 2011, Griffin, Niederhuth et al. 2016, Xu, Wang et al. 2017). Module M6 was also enriched in genes related to the metabolic pathway of anthocyanins, compounds responsible of red colors in plants, but also related to plant stress response (Gould 2004, Landi, Tattini et al. 2015). Anthocyanins seem to be regulated by DNA methylation in several plants, mostly due to overlap between a transposable element and the promoter of *MYB* family transcription factors (Wang, Wang et al. 2020, Zhu, Zhang et al. 2020, Li, Yu et al. 2022). Given that transposable elements are heavily methylated in plants to repress their expression while sometimes overlapping with promoter regions (Ito, Gaubert et al. 2011, Matzke and Mosher 2014), finding co-expression between transposons and anthocyanin signaling pathway genes may indicate that DNA methylated transposable elements play a potential role in anthocyanin gene expression regulation. Our results suggest that those changes would be relevant not only above ground (e.g., fruit maturation, leaf abiotic stress and herbivory responses) but also in roots, where they could respond to defense-growth trade-offs.

Given *E. cicutarium*’s estimated genome size (765 Mbps, 6 fold bigger than *A. thaliana*) and its possible polyploidy, each replication event should have incorporated higher than expected amounts of 5-azaC during the 48h exposure, thus we expected a fast depletion of 5-azaC within cells and a consequential dissipation of its effects during time. In contrast, adult leaves showed more DEGs induced by 5-azaC compared to juvenile leaves. Moreover, the overlap between DEGs affected by 5-azaC in juvenile and adult leaves was not greater than expected by chance, suggesting that the long term response to DNA demethylation by 5-azaC affects different pathways than the short term response, which is a new finding that should be investigated in other species to better understand long-term responses to stress. Among the differentially expressed genes from adult leaves, we found that 5-azaC induced up-regulation of genes related to seed dormancy inhibition and gibberellic acid signaling pathway (the latter is also a contributor to seed dormancy release (Nonogaki 2014)). Acceleration of flowering in plants exposed to 5-azaC is associated to the DNA demethylation of regulatory elements from genes belonging to the gibberellic acid biosynthetic pathway (Burn, Bagnall et al. 1993, Kondo, Miura et al. 2007, Yari, Roein et al. 2021), which explains why we found genes related to this pathway (usually repressed at this age stage) up-regulated by 5-azacytidine (in adult leaves, but not in juvenile leaves). At least another study found that the transcriptome of plants exposed to 5-azacytidine also shows up-regulation of seed dormancy release genes in *Paeonia suffruticosa* (Zhang, Si et al. 2020).

In conclusion, this study generates insights on the effect of DNA methylation inhibition through 5-azaC in gene expression across different tissues and developmental stages in the non-model plant *E. cicutarium*. Mostly expected gene expression patterns were found, such as 5-azaC induced up-regulation of *CMT1, DRM2*, transposable elements and seed dormancy inhibition genes. Nevertheless, unexpected patterns also occurred, such as the magnitude of the 5-azaC effect in the transcriptome of adult leaves compared to juveniles. By characterizing both the transcriptomic response to 5-azaC and the DNA methyltransferase gene set from *E. cicutarium*, this study expands the available genomic resources for non-model species, a need to move forwards current understanding of the epigenetic contribution for adaptation in wild plant populations. In future studies, tracking down plant phenotype changes after 5-azaC treatment can be extended by the study of the transcriptome and methylome, which allows to view the integrative response of plants to 5-azaC, potentially enhancing the identification of which phenotype changes are caused by changes in DNA methylation or solely by gene expression changes. Given the impact of 5-azaC seed treatment in adult/reproductive plants, such experiments could be performed at a longer term range.

## Data availability

The *E. cicutarium* genomic data is publicly available at NCBI genome repository under genome project accession JBDFRG000000000, with the raw reads hosted at NCBI sequence read archive (SRA) repository under accession number SRR28842229. Protein sequences used in the phylogenetic analysis are available at **Supplemental Information** in this journal. RNA-seq raw reads are available at NCBI SRA repository under accession numbers SRR29751957-SRR29751979. Scripts to analyze the data are publicly available at GitHub (https://github.com/rmblazquez/erodium-cicutarium-Transcriptomics-ebd).

## Acknowledgements

We thank Pilar Bazaga and Javier Puy for the assistance with reagents preparation and greenhouse work. We thank AllGenetics for the *E. cicutarium* genome assembly. We thank Summer Blanco and James Leebens-Mack for sharing the protein sequences from their *G. maculatum* genome project and providing input on the draft version.

## Funding

This study was supported by the Spanish Ministerio de Ciencia, Innovación y Universidades, through the “Proyectos de I+D+i” program, with references PID2019-104365GB-I00 and

PID2022-141530NB-C22. We acknowledge support of the publication fee by the CSIC Open Access Publication Support Initiative through its Unit of Information Resources for Research (URICI).

